# Chitosan reduces naturally occurring plant pathogenic fungi and increases nematophagous fungus *Purpureocillium* under field soil conditions

**DOI:** 10.1101/2024.07.01.600678

**Authors:** Raquel Lopez-Nuñez, Jorge Prieto-Rubio, Inmaculada Bautista-Carrascosa, Antonio L. Lidón-Cerezuela, Miguel Valverde-Urrea, Federico Lopez-Moya, Luis V. Lopez-Llorca

## Abstract

Chitosan reduced soil pH, conductivity (CE) and cation exchange capacity (CEC) in pots when applied at field capacity. However, chitosan did not affect these soil physicochemical properties, when applied monthly to agricultural fields. Chitosan did not affect field respiration. Increases in field soil respiration found in chitosan plots, especially in spring-midsummer, were not significant. Although, no differences in soil mineral nitrogen were found, chitosan influenced field soil microbiota. Metabarcoding showed chitosan significantly modifies fungal genera composition of ecologically managed field soil. On the contrary, chitosan caused no significant differences in bacterial taxa composition of field soil. Chitosan coacervates increase naturally occurring nematophagous fungus *Purpureocillium* in soil respect to chitosan solution treated soil and untreated controls. Besides chitosan reduces inoculum of plant pathogenic fungi *Alternaria* and *Fusarium* in field soil. Soil microbial co-occurrence network analysis clustering coefficient (CC) for ITS+V1-V2 regions show that the nematophagous fungus *Pochonia* promoted network clustering into modules. In addition, CC in ITS+V3-V4 regions show that the nematode trapping-fungus *Orbilia* and bacteria belonging to *Acidimicrobiales* and *Cytophagales* also significantly contributed to microbial network clustering in field soil. Our results show that chitosan coacervates increase soil nematophagous microbiota and that both nematode egg-parasites and trapping-fungi help to structure soil microbiota.

## 1. Introduction

The use of chemical pesticides, imposed by demographic changes, is the most common strategy to improve agricultural productivity. However, there is a trend towards the use of ecological additives, such as chitosan, with low environmental impact, instead of chemical synthesis agrochemicals such as nematicides (Bautista-Baños et al., 2005; Lopez-Nuñez et al., 2022). Chitosan is also a source of nitrogen for stimulating plant growth (Pichyangkura and Chadchawanb, 2015). The behaviour of chitosan in soil is related to its cationic nature. This allows electrical interactions with the negatively charged surfaces of clay minerals, modifying its behaviour in soil (Hataf et al., 2018).

Chitosan can modify some soil properties (Reddy et al., 2018). This biopolymer can act as a cohesive agent for clay particles (Hataf et al., 2018). Arid soils are often low on natural polysaccharides which stabilise soil structure (Orts et al., 2000). Chitosan can bind metal ions and limit their leachability, even in the presence of K^+^, Cl^-^ and NO_3_^-^, the dominant ions in soil (Kamari et al., 2011). Furthermore, it can reduce the bioavailability of nickel (Turan, 2019; Heidari et al., 2020) and immobilises chromium when combined with other adsorbents (Najafi et al., 2021). Chitosan is a source of nitrogen, promoting plant growth (Pichyangkura and Chadchawanb, 2015). Chitosan is also an elicitor of plant defences that can trigger physiological and structural responses in the plant, inducing jasmonic acid (JA) and salicylic acid (SA) production (Lopez-Moya et al., 2019; Suarez-Fernandez et al., 2020). Chitosan is active against plant pathogenic nematodes (Khalil and Badawy, 2012), has antiviral and antifungal activity and induces tolerance to abiotic and biotic stresses in several horticultural crops (Iriti and Varoni, 2015; Malerba and Cerana, 2016).

Chitosan sensitivity of filamentous fungi and yeasts increases with carbon and nitrogen limitation (Lopez-Moya et al., 2015). Chitosan permeabilises the membrane of the fungus *Neurospora crassa*, in an energy dependent manner. Conidia are most sensitive to chitosan membrane permeabilization followed by germlings and vegetative hyphae. Therefore, chitosan causes conidial lysis and death within minutes (Palma-Guerrero et al., 2009). Membrane fluidity is a key factor in fungal sensitivity to chitosan (Palma-Guerrero et al., 2010a; Zavala-González et al., 2016). Chitosan-sensitive fungi such as important plant pathogens (eg. *Fusarium* spp. and *Alternaria* spp.) have a high content of polyunsaturated fatty acids (Ren et al., 2021; Chen et al., 2014). In contrast, chitosan-resistant fungi such as nematophagous (*Pochonia chlamydosporia)* or entomopathogens (*Beauveria bassiana),* have a lower presence of polyunsaturated fatty acids in membrane lipids. These fungi express, upon exposure to chitosan, extracellular hydrolytic enzymes (chitosanases, chitinases, proteases) involved in nematode egg-penetration. Furthermore, chitosan increases conidiation in nematophagous and entomopathogenic fungi (Palma-Guerrero et al., 2010b; Palma-Guerrero et al., 2010c).

In this work we study the effect of chitosan solutions or coacervates on soil physicochemical properties in laboratory-controlled conditions at constant humidity (field capacity) and in the field (under conventional and ecological regimes). We also test the effect of chitosan on ecological agriculture soil microbiota using metabarcoding. We also evaluate, in this soil, the effect of chitosan on fungi and bacteria co-occurrence networks.

## 2. Materials and Methods

### 2.1. Chitosan solutions and coacervates

Chitosan powder (Marine BioProducts GmbH, Germany) was dissolved in 0.25M HCl to obtain an initial concentration of 10 mg/mL and pH was adjusted to 5.6. The resulting solution was then dialysed against distilled water for two days and autoclaved. Chitosan solutions were stored at 4°C until used for a maximum of 30 days. Control solutions were prepared likewise but without adding chitosan.

Chitosan was dissolved in sodium acetate buffer (pH 5) to obtain a 3% solution. Chitosan coacervates (CC) were formed by dropping a 3% chitosan solution into 10% sodium hydroxide using a plastic syringe (Terumo Europe NV), with a 0.2 mm diameter outlet. CC were left for 5 minutes in the sodium hydroxide solution. CC were then washed in sterile distilled water to reach pH 8. CC were dried onto sterile filter paper in a laminar flow hood (TELSTAR BV-100) for 24 h. CC were then stored at room temperature in sterile containers.

### 2.2. Application of chitosan to agricultural field soil

Persimmon fields in Pedralba (Valencia, E, Spain), conventionally (39° 35’ 55.25’’ N, 0° 43’ 47.31 W) and ecologically (39° 35’ 52.47’’ N, 0° 43’ 41.47 W) farmed were selected for experiments (Table 1). Soil properties were determined air dried soil sieved through 2mm. Soil pH was measured in a 1:2.5 (w/v) aqueous solution using a pH meter (2001, Crison, Barcelona, Spain). Electrical conductivity was determined in a 1:5 (w/v) aqueous solution using a conductivity meter (model, Crison). The carbonate content was determined using a Bernard calcimeter. Soil organic matter (OM) was determined by wet oxidation using Walkley-Black titration method (Walkley and Black, 1934). Soil texture was determined by the Bouyoucos method (Boyoucus, 1927). Surface soil from both plots was taken for the incubation experiment with pots in growth chambers. Also, these plots were used for an experiment of chitosan application in the field where two treatments were selected: coacervates, only one application at the beginning of the experiment and soluble chitosan applied monthly with the dose divided between the number of applications.

**Table 1:** Physicochemical characteristics of the soils used in this study.

| Type soil | Texture | AD (g/cm<sup>3</sup>) | pH (H<sub>2</sub>O) | CE 1:5 (dS/m) | CaCO<sub>3</sub> (%) | OM (%) |
| --- | --- | --- | --- | --- | --- | --- |
| A Cq E | Sandy Loam | 1.161 | 7.76 ± 0.03 | 0.40 ± 0.03 | 43.18 ± 0.12 | 13.05 ± 1.29 |
| A Cq C | Loam | 1.303 | 8.16 ± 0.08 | 0.20 ± 0.06 | 37.04 ± 1.82 | 3.42 ± 0.05 |

1.BD: Bulk density; CE 1:5: electrical conductivity extracts 1:5; OM: organic matter. A CQ E: Pedralba persimmon ecological; A CQ C: Pedralba Persimmon Conventional. (Lull et al., 2021)

#### 2.2.1 Field experiments

Three 1 x 1 m plots were marked in each field (Fig. 1). Each plot was subdivided into six 33 x 50 cm subplots. Three subplots per plot were randomly selected for treatments. These included: Control (C), 1 mg/mL Chitosan solution (T8L) and Chitosan coacervates (T8C). Selected T8C subplots were treated with chitosan coacervates (9 g/subplot) at the start of the experiment. C and T8L subplots were irrigated monthly (1L/subplot) for nine months with either distilled water (C and T8C) or 1 mg/mL chitosan (T8L). Field soil temperature and electrical conductivity were measured monthly (for nine months) using a WET-2 sensor (HH2 Moisture Meter, Delta-T Devices, Burwell, UK). Respiration rate and CO_2_ concentration were also measured monthly (for nine months) using a EGM-4 environmental gas monitor device (PP System Company, Amesbury, MA, USA). At the end of the experiment, four core samples were collected from each treated subplot with a cylindrical auger (5.35 cm diameter and 12.77 cm long). Soil cores were placed in 15 x 20 cm sterile airtight bags. Soil subsamples (10g) were sieved through a 2mm mesh, then were air dried to measure cation exchange capacity, pH, soil moisture (see below) and mineral nitrogen. Soil mineral nitrogen (nitrate and ammonium) was extracted in 2M KCl and analysed colorimetrically by flow injection (FIAStar 5000, Foss Tecator, Höganäs, Sweden) (Rhoades, 1982). Cation exchange capacity was determined by the sodium acetate sodium chloride method (Rhoades, 1982).

**Figure 1:**
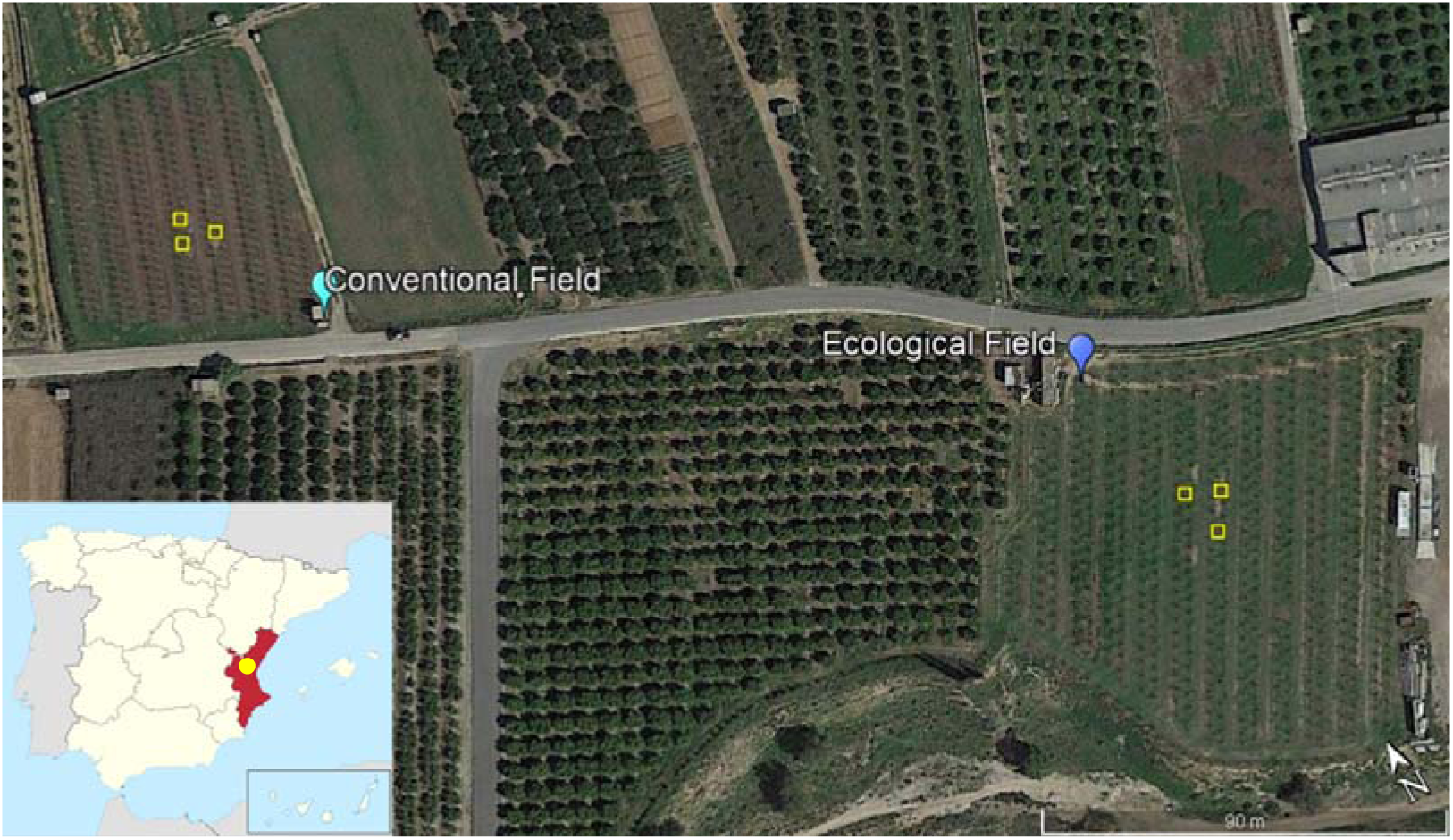
Persimmon experimental fields. Fields were in Pedralba, Valencian Community (East, Spain). Conventional Field received mineral fertilization and usual agronomic practices. Ecological Field received organic fertilization only and no agrochemicals were applied. Experimental plots (1×1 m) where treatments were applied, and soil samples collected are marked by yellow boxes.

#### 2.2.2. Laboratory Trials

Polystyrene cups (200 mL) with a hole in the base covered with glass wool were filled with soil. Cups were incubated in a growth chamber (SANYO, MLR-351H) at 24°C, 60% relative humidity under 16 h light/ 8 h dark photoperiod. Cups with soil were irrigated periodically to maintain field capacity (see below). Ten replicate pots were set per soil (conventional and ecological management) and treatment (as for field experiment).

After 30 days soil from three pots per soil/treatment was pooled and homogenised per triplicate (nine pots sampled). Then soil humidity, pH, electrical conductivity and cation exchange capacity were analysed. This experiment was carried out in duplicate.

### 2.3. Field capacity water content

We placed 12.5 cm of soil in a 15.5 cm long and 3.5 cm wide percolation tube. Water was then added to wet the first 5 cm of soil. The top of the tube was capped with parafilm and aluminium foil, leaving the tap open for 48-72 hours. We then discarded the first centimetre of soil, took a sample of moist soil, and weighed it. We dried the soil at 50 °C to constant weight. We calculated soil field capacity with the formula (Llorca-Llorca, 1991):

Soil Field capacity = (Moist Soil Weight – Dry Soil Weight)/ Dry Humid Weight

### 2.4. Physico-chemical analysis of soils

Soil moisture, conductivity/salinity, pH, texture, and cation exchange capacity were determined for all soil samples collected (Llorca-Llorca, 1991). Three measurements were taken per each physicochemical parameter for treatment and soil type.

### 2.5. Soil metabarcoding

Soil samples from ecological orchard plots were taken from each subplot and treatment as for physicochemical determinations (see before) at the end of the experiment. On the same day of collection, DNA was extracted from fresh soil (250 mg per soil sample), using DNeasy PowerSoil Pro Kit (Qiagen, Germantown, MD, USA). DNA samples were sent to Macrogen Inc. (Seoul, Korea), where they were amplified and using specific fungi (ITS) and bacteria (V1-V2, V3-V4) primers (Table 2) and sequenced by the Illumina Miseq platform using v3 reagent kit. DNA reads obtained were analysed using OmicsBox 3.0 package to identify the microorganisms present in soil samples.

**Table 2:**
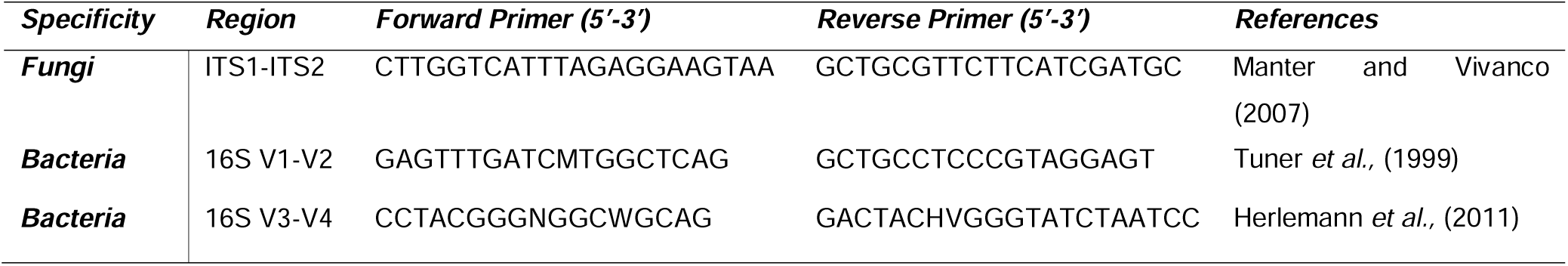
Primers used in this study.

### 2.6. Soil microbe co-occurrence networks

Fungal and bacterial communities characterized from ITS, V1-V2 and V3-V4 amplicon sequencing were analysed through co-occurrence networks by using the SParse InversE Covariance Estimation for Ecological Association Inference (SPIEC.EASI) pipeline in R package (Kurtz et al., 2015). This network-based approach allowed to frame both fungal and bacterial communities into a same co-occurrence network (Wagg et al., 2019). Before network inferring, OTUs that occurred > 1 % and more than five samples were maintained in the datasets and re-scaled to the proportion of the minimum sequencing depth (32,672 reads for fungi in the ITS dataset, 38,709 for bacteria in the V1-V2 and 38,011 for bacteria in the ITS+V3-V4). The inference was carried out by combining the amplicon pair dataset, ITS+V1-V2 and ITS+V3-V4. We fitted the spiec.easi function with Meinshausen-Buhlmann’s neighbourhood selection method and the lambda minimum ratio at 0.01. From the spiec.easi object, we extracted the OTU adjacency matrix with the symBeta function to infer the network graphs and network properties of OTUS from the Gephi software (Bastian et al., 2009). In particular, we determined the degree centrality, that counts the number of links per OTU and is weighted by the occurrence frequency per linked OTU pairs (Gouveia et al., 2021), the modularity class for each OTU embedded in the network, i.e. the module which an OTU belong to, and the clustering coefficient, that measures the extent of an OTU to cluster with others into a module (Latapy 2008).

### 2.7. Statistical analysis

Results from pot tests were analysed with a Three-Way ANOVA to determine statistical differences for each variable tested (pH, Conductivity, Cation Exchange Capacity), with the factors soil, treatment (fixed and orthogonal) and experiment (random and orthogonal) at the end of the experiment (30 days).

For the field test variables (pH, Conductivity, Cation Exchange Capacity and Mineral Nitrogen), a Two-Way ANOVA of soil and treatment (fixed and orthogonal) was performed for the last data collection time of the field experiment (9 months).

Then, a Three-Way ANOVA was performed to analyse the differences of each variable (Respiration Rate, Conductivity, Moisture and Temperature), with the factors soil, treatment and time (fixed and orthogonal). The ANOVA requirements were tested with the DHARMa package (Hartig 2022).

For the ecological soil Metabarcoding analysis, the OmicsBox 3.0 program was used to obtain relative abundances of Phylum, Order, Genus, and Species for the ITS, V1-V2 and V3-V4 primers, with the Kraken 2.1.2 function (Wood et al., 2019; Wood and Salzberg, 2014). Abundances above 1% (relative abundance), were taken for statistical analyses. The mean relative abundance and standard error were calculated with Excel.

To study the differences of Phylum, Genus, Order and Species present in the ecological soil according to treatment, a multivariate generalised linear model (GLM) with a Gaussian distribution of the error (“manyglm” function in the “mvabund” package) was performed. A univariate GLM with a Gaussian family error distribution was then performed for each variable to analyse the differences between abundances in Genera and Species for ITS primers. Treatment was considered as a predictor variable in the analysis. We conducted pairwise comparisons with estimated marginal means (“emmeans” function and package; Lenth, 2023) using Sidak’s HSD test for GLM data.

The effect of taxonomy on network metrics was assessed by fitting linear regression models for each amplicon pair data set, ITS+V1-V2 and ITS+V3-V4. A t-test was performed on the estimated values to detect taxa that significantly explained the results of the network metrics.

All statistical analyses were performed with R software (version 4.2.2) (R Core Team, 2023).

## 3. Results

### 3.1. Chitosan reduces potted soil pH conductivity and cation exchange capacity

Chitosan solutions significantly reduced soil pH (ANOVA; p value=0.001) (Fig. 2A; Table S1) and conductivity (CE) (ANOVA; p value=0.04) (Fig. 2C, Table S3) when water content is maintained at field capacity in the pot experiment. Both chitosan solutions and coacervates reduced soil cation exchange capacity (CEC) respect to controls in the pot experiment (ANOVA; p value=0.03) (Fig 2E, Table S5). However, under field conditions when applied monthly, chitosan did not alter field soil pH, CE, and CEC (ANOVA; p value = 0.5, p value= 0.3, p value= 0.1; Fig. 2B, D, F; Tables S2, S4, S6). In the field experiment soil CE was lower for March-July than for November-February recordings for both soil managements (Fig. S1). In June and July in the organic soil, conductivity could not be recorded because of low soil humidity for high temperatures and low rainfall (Fig. S2, S3). Chitosan application to field soil had no significant effect on soil mineral nitrogen content (Fig. S4).

**Figure 2:**
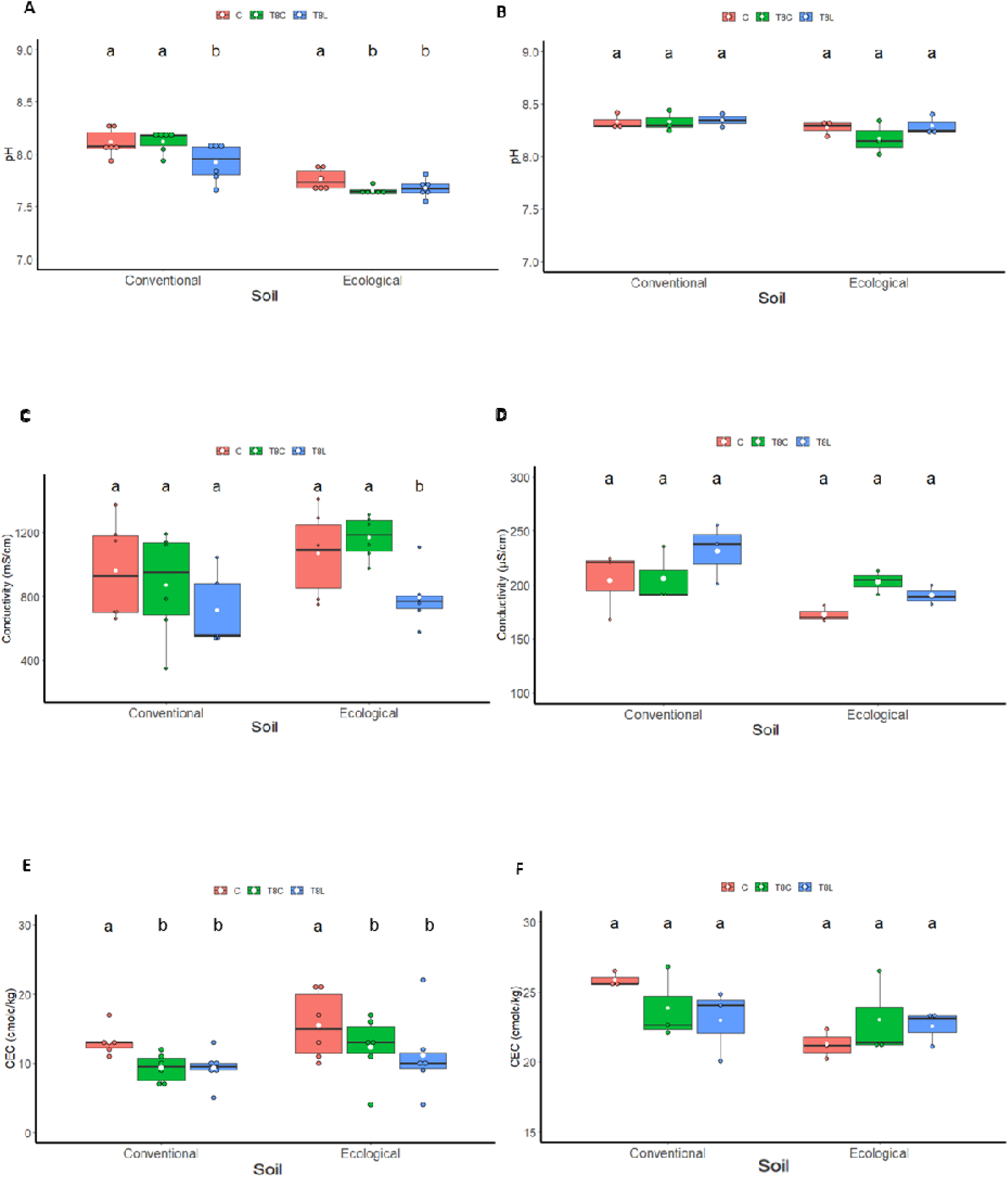
Effect of chitosan on soil chemical properties: soil pH A), B), conductivity C), D) and cation exchange capacity E), F). Treatments: Control (C, untreated), chitosan coacervates (T8C) and chitosan solution (T8L). Experiments: Growth chambers (A, C, E), Field (B, D, F). Lower case letters indicate significant differences between treatments for each soil type.

### 3.2. Chitosan does not affect field soil respiration under field conditions

A trend of increased respiration was observed in the chitosan treatments mainly from March to June for the conventional orchard soil (Fig. 3A) and from March and May for the ecological orchard soil (Fig. 3B). However, these differences were not significant. Irrespective of treatments, field soil respiration significantly (ANOVA; p-value > 0.001, Table S7) increased in both management regimes (conventional and ecological) from March until July. This period corresponds with a steady significant increase in soil temperature for both conventional and ecological regimes (Fig. S2 A, B). Soil moisture increased in the march recording (Fig S2 C, D). This corresponded, in turn, with an increase in precipitation (Fig. S3).

**Figure 3:**
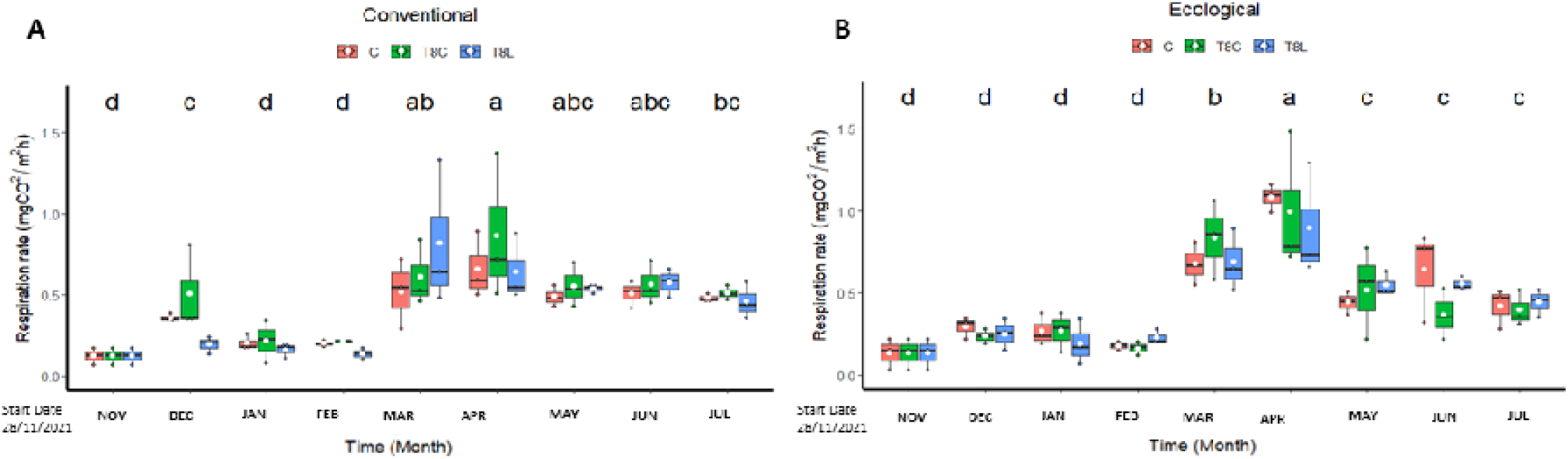
Effect of chitosan on field soil respiration. Soil was under conventional (A), or ecological (B) regimes Treatments: Field (C, untreated), chitosan coacervates (T8C) and chitosan solution (T8L). Low-case letters show significant differences between the different times. Level of significant differences p-value<0.05.

### 3.3. Chitosan modifies soil mycobiota

Chitosan significantly (multivariate GLM, p.value 0.001, Table S8) modifies fungal genera composition of ecological field soil (Fig. 4A). Conversely, chitosan caused no significant differences in bacterial taxa composition of the same soil respect to untreated controls (multivariate GLM, p value > 0.001; Figs. 5B).

**Figure 4:**
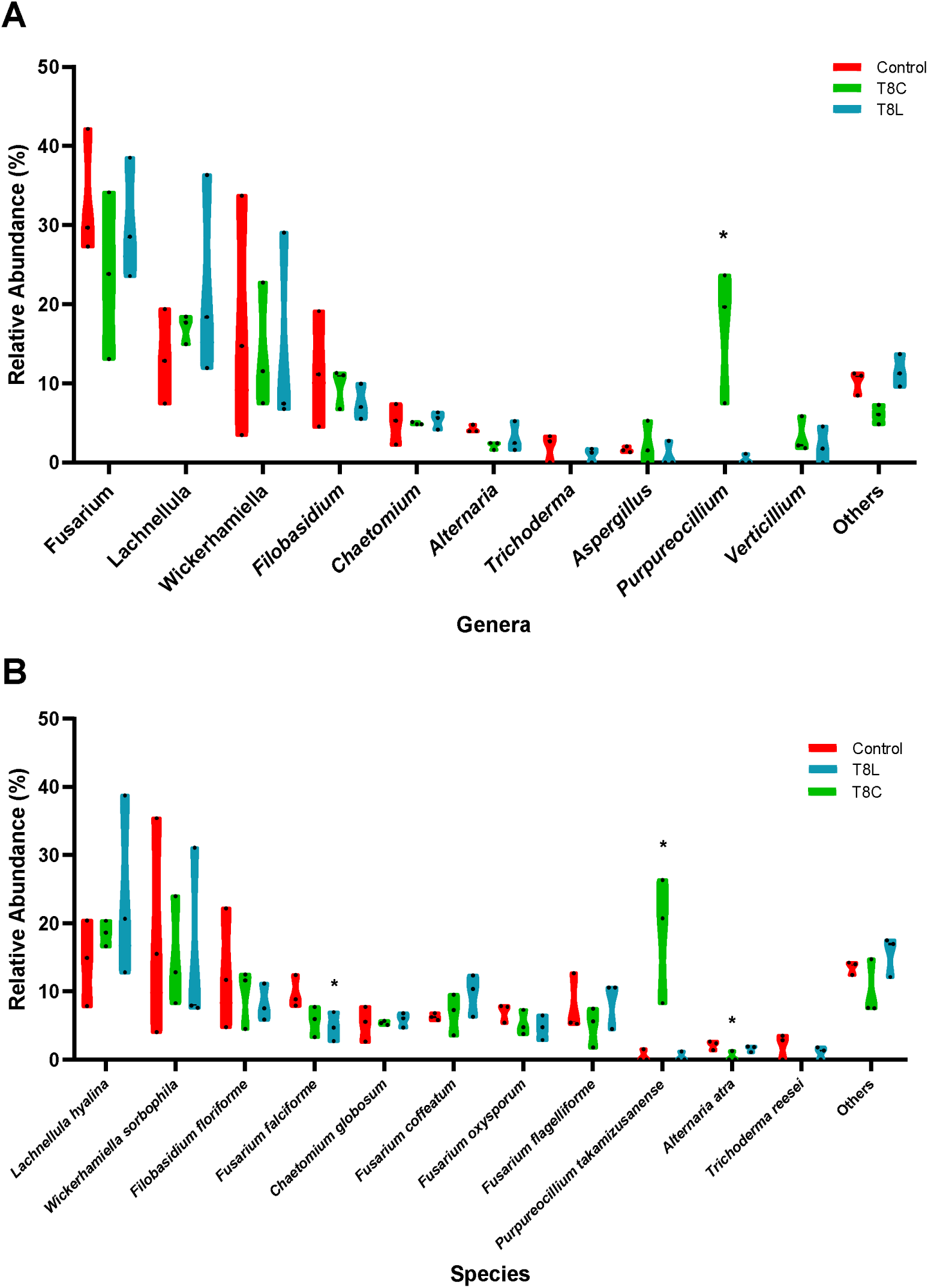
Relative abundance (%). Asterisks mark significant differences (p value > 0.005), of the treatments with respect to the control for each genera (A) and species (B).

**Figure 5:**
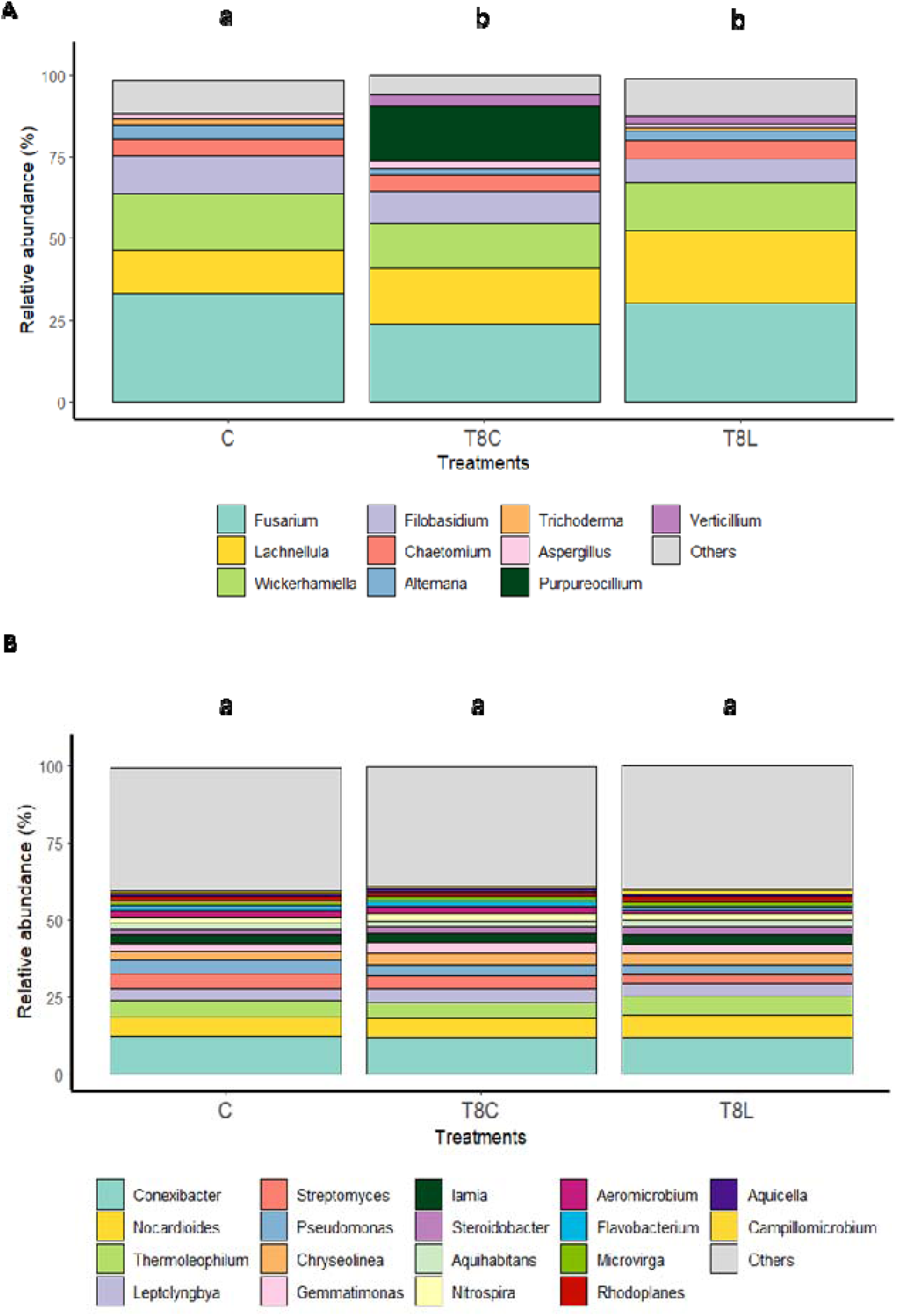
Effect of chitosan on field soil microbiota. A) Fungal genera (ITS primers) and B) bacterial genera (V1-V2 primers). Treatments: Field (C, untreated), chitosan coacervates (T8C) and chitosan solution (T8L). Different letters indicate significant differences (p-value < 0.05).

### 3.4. Chitosan reduces naturally occurring plant pathogenic fungi in field soil

*Fusarium* was the fungal genus most present (33-23%) in field samples (Fig 5A), followed by *Lachnellula (22-13%)*, *Wickerhamiella* (17-14%) and *Filobasidium* (11-7%) (Table S9). Other genera, including *Alternaria*, showed 5% or less relative abundance (Fig.5A). Chitosan coacervates tended to reduce the relative abundance of *Fusarium* and *Alternaria*, although no significant differences were found. Presence of the plant pathogenic species *Fusarium falciforme* (50% reduction, Table S10), in soil was significantly reduced (univariate GLM, p.value= 0.03, Table S11), by chitosan solution (Fig. 4B). Chitosan coacervates significantly reduced (univariate GLM p.value = 0.01, Table S11) the relative abundance of the phytopathogenic species *Alternaria atra* (20% reduction, Table S10), respect to untreated controls.

### 3.5. Chitosan coacervates increase naturally occurring nematophagous fungus *Purpureocillium* in field soil

Chitosan coacervates significantly (univariant GLM, p value = 0.006, Table S12), increased (*ca.* 4500%), naturally occurring nematophagous fungus *Purpureocillium* in field soil, (Fig. 5A). Significant differences were found for the variable fungal species (multivariate GLM, p.value = 0.044; Fig. S5, Table S13) between control and chitosan coacervate treatments. Chitosan coacervates significantly increase (*ca.* 3500%) the presence of the invertebrate pathogen *Pupureocillium takamizusanense* in soil (univariate GLM, p =0.006, Fig.5B, Table S11).

### 3.6. Nematophagous fungi structure soil microbiota

The use of ITS+V1-V2 and ITS+V3-V4 regions revealed variations in the co-occurrence network outcomes (Figure 6; Tables S14 and S15). However, we show that the weighted degree centrality (WDC) parameter could not allow to detect contrasting influence of microbial groups within the network of each amplified region in the ITS+V3-V4 subset, only marginally detected in bacteria that belonged to *Acidimicrobiales*, (Figure 6A).

**Figure 6:**
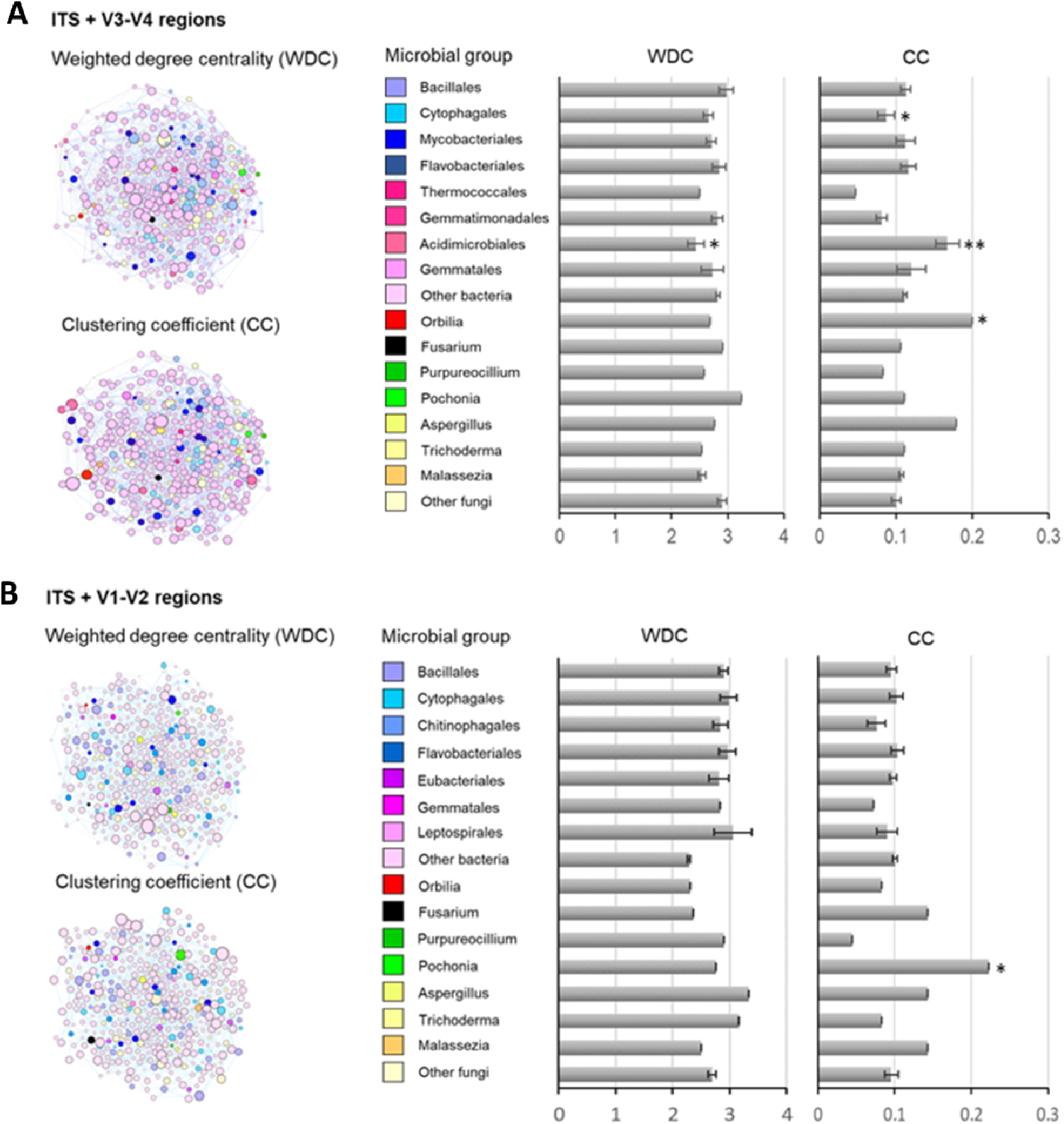
Microbial co-occurrence networks by amplified region pairs, ITS (fungi) and V1-V2/V3-V4 (bacteria) in soil ecological. Networks were inferred from SPIEC.EASI R package. Graphs and network metrics -weighted degree centrality (WDC), i.e. the interaction frequency between OTU pairs, and clustering coefficient (CC), i.e. the extent of an OTU to cluster with others into modules-, were determined in Gephi software. WDC and CC values are also shown in bar graphs by microbial group, bacteria and fungi at order and genus taxonomic levels, respectively. The effect of microbial taxa identity on the network metrics were evaluated by linear models and t-tested, and those significantly explaining WDC and CC are marked by the significance level: ‘**’ p < 0.01, ‘*’ p < 0.05.

Co-occurrence network (CC) parameter for ITS+V1-V2 regions showed that the nematode egg-parasitic fungi *Pochonia* (CC = 0.22) promoting network clustering into modules (number of modules = 15) (Figure 6B; Table S16). Evaluating the cooccurrence type we show *Pochonia chlamydosporia* with a positively interaction to xylan degrading (*Humisphaera*), N-fixing (*Leptolyngbya*) and sulphate-reducing bacteria (*Rubrobacter*) (Table S18). Therefore, we have detected antagonistic interactions with soil bacteria such as *Aquihabitans* spp., a gram-negative bacteria, *Lectolyngbya* spp., a world-wide distributed cyanobacteria and *Proteatibacter* spp., a widely distributed soil bacteria (Table S18). The ITS+V3-V4 regions showed that the nematode-trapping fungi *Orbilia* (CC = 0.20) and *Acidimicrobiales* (CC = 0.17 ± 0.02) and *Cytophagales* (CC = 0.04 ± 0.01) significantly contributed to network clustering into modules (n = 29) (Figure 5B; Table S17). *Orbilia oligospora,* show synergistic cooccurrence with a wide group of soil bacteria (*Nakamurella* spp, *Nocardioides* spp. or *Vulgatibacter* spp.). By the other side, *O. oligospora* show a competitive behaviour with important soil borne fungal pathogens like *Talaromyces* spp. and *Aspergillus* spp. species (Table S19).

*Acidimicrobium ferrooxidans,* an extremophile bacterial able to growth under extremely low pH conditions (pH<2), shows positive interactions with soil-living bacteria such as *Massilia* spp., *Nitrospira* spp. or *Stella* spp. However, this bacterium has antagonistic effect on *Jiangella* spp. *Hymenobacter* spp. and *Limnoglobus* spp. bacteria present in crop soils. Inside of the *Cytophagales*, we have shown *Cytophaga hutchinsonii* playing a key role in the microbial community. This bacterium shows positive interactions with many soil-born bacteria (*Calothrix* spp., *Chitinophaga* spp. or *Lysobater* spp.). Interestingly, *C. hutchinsonii* play a competitive role against important soil fungal pathogens like *Fusarium oxysporum* and *Verticillium dhalie*.

## 4. Discussion

Chitosan applied maintaining soil water content at field capacity in pots for a month, significantly reduced soil pH, CE and CEC. The slight reduction of pH in the soil induced by chitosan could be simply due to the weak acidity of chitosan solutions. This effect was not found in field soil. This was perhaps due to lower volumes of these chitosan solutions applied monthly. The high calcium carbonate content of both soils tested may have neutralised the chitosan solutions. The reduction of soil CE by chitosan in pots could be due to the mopping capacity of chitosan (polycation) of ions present in the soil solution (Kamari et al., 2011). Chitosan solutions and coacervates reduced soil cation exchange capacity (CEC) respect to controls for potted soils. When applied to sodium montmorillonite, chitosan intercalates in the layers of the clay (Darder et al., 2003) both reducing the negative charges for cation exchange and immobilizing chitosan in soils. In our pot study, by doing this chitosan may have displaced exchangeable cations from the clay complex, thus reducing CEC. However, this was not found when chitosan was applied monthly in the field. The regime of chitosan irrigation (field capacity vs. monthly applications) could account for a lower chitosan presence in field soil than in the pots. This may have made the chitosan displacement of cations of the clay complex in field soil less efficient than in pots. Taken together our results suggest that chitosan can be applied to agricultural fields without affecting CEC, a soil fertility related key parameter (Anderson et al., 2023)

Undissolved chitosan added to soil (5%w/w) caused N increase (Ammonium and nitrogen), respect to untreated controls in a microcosm experiment (Sawaguchi et al., 2015). In our study chitosan application to field soil had no significant effect on soil mineral nitrogen content due mainly to the high mobility of mineral in soils. Our treatments also involved less chitosan applied to soil than in the microcosm. This and the time lapse (9 months) for N soil content testing may explain our results. In soil incubation experiments with chitosan, soil respiration was found to increase with chitosan concentration (Nkoh et al., 2024). In our field study, chitosan treatments resulted in increases in soil respiration, especially in spring-midsummer. This effect, although no significant, could be due to sudden organic N input when chitosan was added to our microplots.

Our metabarcoding study shows that chitosan significantly modifies fungal genera composition of ecological field soil. Chitosan coacervates increase naturally occurring nematophagous fungus *Purpureocillium* in soil respect to chitosan solution treated soil and untreated controls. Chitosan increases ca. 6000%, conidiation of *Purpureocillium* (Palma-Guerrero et al., 2010c) cultures respect to control media with no chitosan. The similar (ca. 4000%) increase in the relative abundance of *Purpureocillium* found in this work could be due to chitosan induction of conidiation of the fungus naturally occurring in soil. The highly sporulating capacity of this chitosan tolerant fungus (Gortari and Hours, 2016) could explain our results. *Purpureocillium lilacinum* has been applied combined with chitosan for managing root knot nematodes (Giannakou et al, 2020; Zhan et al., 2021). The species *Purpureocillium takamizusanense* identified in our study, has been isolated as an entomopathogenic fungus (Nguyen et al., 2022). Future studies should evaluate the effect of chitosan on the performance of the fungus in the field for insect/nematode pest biomanagement. These studies could include augmentative natural biocontrol and enhanced biocontrol with inundative or sustained additions of inoculum of the fungus. Chitosan particles should be used in these studies, since chitosan solutions did not enhance naturally occurring *Pupureocillium* on soil.

We also found that the abundance of *Alternaria atra* and *Fusarium falciforme* decreased in soil treated with a chitosan solution. These two fungal species cause diseases in several crops worldwide (Bonthala et al., 2021; Trolinger et al., 2017). Therefore, soil treatment with chitosan could be a sustainable alternative for managing these fungal plant pathogens. Furthermore, our co-occurrence network analyses (see below) show *Purpureocillium* negatively related to *Alternaria atra* and *A. rosae* (Table S18). *Purpureocillium* is a fungus well known to produce antimicrobial secondary metabolites (Chen and Hu, 2021). Future studies should investigate the mechanisms involved in the antagonism of *Purpureocillium* spp. to *Alternaria* spp. in soil

Metagenomics on soil exposed to chitin-rich exoskeletons has been a source of gene sequences encoding chitin-chitosan degrading enzymes (Li et al., 2015; Stöveken et al., 2015). Most of these sequences were from bacterial origin. Our metabarcoding analysis shows that chitosan application during 9 months to field soil do not change bacterial taxa profiles. Perhaps time of exposure to chitin/chitosan could account for these differences.

We have carried out a microbial diversity and ecological network analysis (Barberán et al., 2012). Our results show that the two main ecological groups of nematode destroying fungi (Barron 1997): nematode trapping (*Orbilia* spp.) and egg parasites (*Pochonia* spp.), promote soil microbe network clustering into modules. Nematophagous fungi interact with nematodes, the most abundant animal taxon in soil (Dervash et al., 2018). Most soil nematodes are bacterivorous (De Mesel et al., 2004), so nematophagous fungi are related to soil bacteria too. Our co-occurrence network analyses show the nematode egg parasite fungus *Pochonia* positively related to xylan degrading (*Humisphaera*), N-fixing (*Leptolyngbya*) and sulphate-reducing bacteria (*Rubrobacter*). These soil prokaryotes could help with nutrient acquisition by the fungus. *Nocardioides*, a hydrocarbon degrader, antibiofilm and antibiotic producer filamentous bacterium is negatively correlated with *Pochonia* and positively with *Orbilia*. This and other bacteria (*Paraflavitalea*, Chitinophagaceae) also positively related with *Orbilia*, can degrade chitin in soil. Root nodule bacteria (*Microvirga* and *Botea*) are positively correlated with the nematode trapping fungus. *Pochonia* can be endophytic in crop plants (Manzanilla-Lopez et al., 2013). This can be beneficial for plants defence of these against soil-borne pathogens. Nematode egg fungal parasites are multitrophic organisms than can be enhanced by chitosan (Escudero et al., 2016; 2017). In this work we show the proof that chitosan application in soil enhances *P. lillacinum* recruitment and the promotion of *P. chlamydosporia* as key fungus to structure microbial communities in soil. Bacteria belonging to *Acidimicrobiales* and *Cytophagales* also significantly contributed to network clustering in field soil. These groups relate to iron redox processes (Garber et al., 2021) and carbohydrate polymer (chitin, pectin, cellulose) turnover (Mohapatra et al., 2022) in soil respectively.

They play a key role recruiting soil-borne bacterial essential to maintain soil health. In addition, we show that *C. hutchinsonii* is an antagonistic microorganism against two important plant pathogenic fungi such as *V. dhaliae* and *F. oxysporum* (Kausar et al., 2021). Future studies could combine the use of these fungi with chitosan to treat diseases in various agricultural crops. Our work opens new and promising possibilities to develop integrated strategies based on the use of chitosan formulations to improve soil health and for managing important plant diseases caused by plant parasitic nematodes.

## Author Contributions

Investigation, methodology, data curation, formal analysis, writing (original draft) and visualization: R.L.-N.; curation data, formal analysis and writing (original draft): R.L.-N., M.V.-U and J.P-R; methodology and sample collection: R.L.-N., I.B.-C, A.L.L.-C. and L.V.L.-L.; supervision, methodology and reviewing: F.L.-M.; supervision, methodology, writing (original draft), reviewing and funding acquisition: L.V.L.-L. All authors have read and agreed to the published version of the manuscript.

## Acknowledgements

We would like to thank C.R.A, Agustin Chumillas Roldán, from the Plant Pathology Laboratory (University of Alicante) for his technical support.

## References

Anderson, A., Khaleel, A., cdutter., Blauwet, M., Flores, A., Miller, B., and Burras, C.L (2023). Introduction to Soil Science. Iowa State University Digital Press Ames, Iowa Consulted: 13/06/2024. (https://iastate.pressbooks.pub/introsoilscience/chapter/cec-aec/)

Barberán, A., Bates, S. T., Casamayor, E. O., and Fierer, N. 2012. Using network analysis to explore co-occurrence patterns in soil microbial communities. The ISME journal, 6(2), 343–351. 10.1038/ismej.2011.119

Barron, G. L. (1977). The nematode-destroying fungi. Canadian Biological Publications Ltd.

Bastian, M., Heymann, S., and Jacomy, M. 2009. Gephi: an open source software for exploring and manipulating networks. In Proceedings of the international AAAI conference on web and social media (Vol. 3, No. 1, pp. 361–362). 10.1609/icwsm.v3i1.13937

Bautista-Baños, S., Hernández-Lauzardo, A. N., del Valle, M. V., Bosquez-Molina, E., and Sánchez-Domínguez, D. 2005. Quitosano: una alternativa natural para reducir microorganismos postcosecha y mantener la vida de anaquel de productos hortofrutícolas. Revista Iberoamericana de tecnología postcosecha, 7(1), 1–6.

Bonthala, B., Small, C. S., Lutz, M. A., Graf, A., Krebs, S., Sepúlveda, G., and Stam, R. 2021. ONT-based draft genome assembly and annotation of *Alternaria atra*. Molecular Plant-Microbe Interactions, 34(7), 870–873. 10.1094/MPMI-01-21-0016-A

Bouyoucos, G. J. 1927. The hydrometer as a new method for the mechanical analysis of soils. Soil science, 23(5), 343–354.

Chen, J., Zou, X., Liu, Q., Wang, F., Feng, W., and Wan, N. 2014. Combination effect of chitosan and methyl jasmonate on controlling *Alternaria alternata* and enhancing activity of cherry tomato fruit defense mechanisms. Crop Protection, 56, 31–36. 10.1016/j.cropro.2013.10.007

Chen, W., and Hu, Q. 2021. Secondary metabolites of *Purpureocillium lilacinum*. Molecules, 27(1), 18.

Darder, M., Colilla, M., and Ruiz-Hitzky, E. 2003. Biopolymer− clay nanocomposites based on chitosan intercalated in montmorillonite. Chemistry of Materials, 15(20), 3774–3780. 10.3390/molecules27010018

De Mesel, I., Derycke, S., Moens, T., Van der Gucht, K., Vincx, M., and Swings, J. 2004. Top-down impact of bacterivorous nematodes on the bacterial community structure: a microcosm study. Environmental Microbiology, 6(7), 733–744. 10.1111/j.1462-2920.2004.00610.x.

Dervash, M., Bhat, R., Mushtaq, N., and Singh, V. 2018. Dynamics and importance of soil mesofauna. International Journal of Advance Research in Science and Engineering, 7(4), 2010–2019.

Escudero, N., Ferreira, S. R., Lopez-Moya, F., Naranjo-Ortiz, M. A., Marin-Ortiz, A. I., Thornton, C. R., and Lopez-Llorca, L. V. 2016. Chitosan enhances parasitism of *Meloidogyne javanica* eggs by the nematophagous fungus *Pochonia chlamydosporia*. Fungal biology, 120(4), 572–585. 10.1016/j.funbio.2015.12.005

Escudero, N., Lopez-Moya, F., Ghahremani, Z., Zavala-Gonzalez, E. A., Alaguero-Cordovilla, A., Ros-Ibañez, C., Lacasa, A., Sorribas, F.J., and Lopez-Llorca, L. V. 2017. Chitosan increases tomato root colonization by *Pochonia chlamydosporia* and their combination reduces root-knot nematode damage. Frontiers in plant science, 8, 1415. 10.3389/fpls.2017.01415

Garber, A. I., Cohen, A. B., Nealson, K. H., Ramírez, G. A., Barco, R. A., Enzingmüller-Bleyl, T. C., Gehringer M. M., and Merino, N. 2021. Metagenomic insights into the microbial iron cycle of subseafloor habitats. Frontiers in Microbiology, 12, 667944. 10.3389/fmicb.2021.667944

Giannakou, I. O., Tasoula, V., Tsafara, P., Varimpopi, M., and Antoniou, P. 2020. Efficacy of *Purpureocillium lilacinum* in combination with chitosan for the control of *Meloidogyne javanica*. Biocontrol Science and Technology, 30(7), 671–684. 10.1080/09583157.2020.1756227

Gortari, M. C., and Hours, R. A. 2016. *Purpureocillium lilacinum* LPSC# 876: Producción de conidias en cultivos sobre sustratos sólidos y evaluación de su actividad sobre *Nacobbus aberrans* en plantas de tomate. Revista de la Facultad de Agronomía, La Plata, 115(2), 239–249.

Gouveia C, Móréh Á, Jordán F. 2021. Combining centrality indices: Maximizing the predictability of keystone species in food webs. Ecological Indicators, 126, 107617. 10.1016/j.ecolind.2021.107617

Hataf, N., Ghadir, P., and Ranjbar, N. 2018. Investigation of soil stabilization using chitosan biopolymer. Journal of Cleaner Production, 170, 1493–1500. 10.1016/j.jclepro.2017.09.256

Hartig F. DHARMa: Residual Diagnostics for Hierarchical (Multi-Level / Mixed) Regression Models 2022.

Heidari, J., Amooaghaie, R., and Kiani, S. 2020. Impact of chitosan on nickel bioavailability in soil, the accumulation and tolerance of nickel in *Calendula tripterocarpa*. International Journal of Phytoremediation, 22(11), 1175–1184. 10.1080/15226514.2020.1748564

Iriti, M., and Varoni, E. M. 2015. Chitosan-induced antiviral activity and innate immunity in plants. Environmental Science and Pollution Research, 22(4), 2935–2944. 10.1007/s11356-014-3571-7

Kamari, A., Pulford, I. D., and Hargreaves, J. S. J. 2011. Chitosan as a potential amendment to remediate metal contaminated soil—A characterisation study. Colloids and Surfaces B: Biointerfaces, 82(1), 71–80. 10.1016/j.colsurfb.2010.08.019

Kausar, R., Iram, S., Ahmad, K. S., and Jaffri, S. B. 2021. Molecular characterization of *Fusarium solani* and *Fusarium oxysporum* phyto-pathogens causing mango maturity malconformation. Archives of Phytopathology and Plant Protection, 54(17-18), 1372–1390. 10.1080/03235408.2021.1910417

Khalil, M. S., and Badawy, M. E. 2012. Nematicidal activity of a biopolymer chitosan at different molecular weights against root-knot nematode, *Meloidogyne incognita*. Plant Protection Science, 48(4), 170–178. 10.17221/46/2011-pps

Kurtz, Z. D., Müller, C. L., Miraldi, E. R., Littman, D. R., Blaser, M. J., and Bonneau, R. A. 2015. Sparse and compositionally robust inference of microbial ecological networks. PLoS computational biology, 11(5), e1004226. 10.1371/journal.pcbi.1004226

Latapy, M. (2008). Main-memory triangle computations for very large (sparse (power-law)) graphs. Theoretical computer science, 407(1-3), 458–473. 10.1016/j.tcs.2008.07.017

Lenth, R. V., Buerkner, P., Herve, M. J., Jung, M., Love, J., Miguez, F., Riebl, H., and Singmann, H. (2023). emmeans: Estimated Marginal Means, aka Least-Squares Means. Version 1.9.0. https://cran.r-project.org/package=emmeans/.

Li, H., Fei, Z., Gong, J., Yang, T., Xu, Z., and Shi, J. 2015. Screening and characterization of a highly active chitosanase based on metagenomic technology. Journal of Molecular Catalysis B: Enzymatic, 111, 29–35. 10.1016/j.molcatb.2014.11.005

Llorca-Llorca, R. 1991. Prácticas de edafología. Universidad Politécnica de Valencia.

Lopez-Moya, F., Colom-Valiente, M. F., Martinez-Peinado, P., Martinez-Lopez, J. E., Puelles, E., Sempere-Ortells, J. M., and Lopez-Llorca, L. V. 2015. Carbon and nitrogen limitation increase chitosan antifungal activity in *Neurospora crassa* and fungal human pathogens. Fungal Biology, 119(2-3), 154–169. 10.1111/aab.12199

Lopez-Moya, F., Suarez-Fernandez, M., and Lopez-Llorca, L. 2019. Molecular Mechanisms of Chitosan Interactions with Fungi and Plants. Int. J. Mol. Sci., 20, 332. 10.3390/ijms20020332

Lopez-Nuñez, R., Suarez-Fernandez, M., Lopez-Moya, F., and Lopez-Llorca, L. V. 2022. Chitosan and nematophagous fungi for sustainable management of nematode pests. Frontiers in Fungal Biology, 3, 980341. 10.3389/ffunb.2022.980341

Malerba, M., and Cerana, R. 2016. Chitosan effects on plant systems. International journal of molecular sciences, 17(7), 996. 10.3390/ijms17070996

Manzanilla-Lopez, R. H., Esteves, I., Finetti-Sialer, M. M., Hirsch, P. R., Ward, E., Devonshire, J., and Hidalgo-Díaz, L. 2013. *Pochonia chlamydosporia*: Advances and challenges to improve its performance as a biological control agent of sedentary endo-parasitic nematodes. Journal of Nematology, 45(1), 1–7. PMID: 23589653; PMCID: PMC3625126.

Najafi, Z., Golchin, A., and Alamdari, P. 2021. Comparison of the efficiency of different chitosan composites in immobilisation of chromium in contaminated soils. Archives of Agronomy and Soil Science, 1–14. 10.1080/03650340.2021.1909720

Nguyen, N. H., Tamura, T., and Shimizu, K. 2022. Draft Genome Sequence of *Purpureocillium takamizusanense*, a Potential Bioinsecticide. Microbiology Resource Announcements, 11(7), e00268–22. 10.1128/mra.00268-22

Nkoh, J. N., Guan, P., Li, J. Y., and Xu, R. K. 2024. Effect of carbon and nitrogen mineralization of chitosan and its composites with hematite/gibbsite on soil acidification of an Ultisol induced by urea. Chemosphere, 349, 140896. 10.1016/j.chemosphere.2023.140896

Mohapatra, M., Manu, S., Dash, S. P., and Rastogi, G. 2022. Seagrasses and local environment control the bacterial community structure and carbon substrate utilization in brackish sediments. Journal of Environmental Management, 314, 115013. 10.1016/j.jenvman.2022.115013

Orts, W. J., Sojka, R. E., and Glenn, G. M. 2000. Biopolymer additives to reduce erosion-induced soil losses during irrigation. Industrial Crops and Products, 11(1), 19–29. 10.1016/S0926-6690(99)00030-8

Palma-Guerrero, J., Huang, I.-C., Jansson, H.-B., Salinas, J., Lopez-Llorca, L. V., and Read, N. D. 2009. Chitosan permeabilizes the plasma membrane and kills cells of *Neurospora crassa* in an energy dependent manner. Fungal Genetics and Biology, 46(8), 585–594. 10.1016/j.fgb.2009.02.010

Palma-Guerrero, J, Lopez-Jimenez, J. A., Pérez-Berná, A. J., Huang, I.-C., Jansson, H.-B., Salinas, J., Villalaín, J., Read, N. D., and Lopez-Llorca, L. V. 2010a. Membrane fluidity determines sensitivity of filamentous fungi to chitosan. Molecular Microbiology, 75(4), 1021–1032. 10.1111/j.1365-2958.2009.07039.x

Palma-Guerrero, Javier, Gómez-Vidal, S., Tikhonov, V. E., Salinas, J., Jansson, H. B., and Lopez-Llorca, L. V. 2010b. Comparative analysis of extracellular proteins from *Pochonia chlamydosporia* grown with chitosan or chitin as main carbon and nitrogen sources. Enzyme and Microbial Technology, 46(7), 568–574. 10.1016/j.enzmictec.2010.02.009

Palma-Guerrero, Javier, Larriba, E., Güerri-Agulló, B., Jansson, H.-B., Salinas, J., and Lopez-Llorca, L. V. 2010c. Chitosan increases conidiation in fungal pathogens of invertebrates. Applied Microbiology and Biotechnology, 87(6), 2237–2245. 10.1007/s00253-010-2693-1

Pichyangkura, R., and Chadchawan, S. 2015. Biostimulant activity of chitosan in horticulture. Scientia Horticulturae, 196, 49–65. 10.1016/j.scienta.2015.09.031

R Core Team (2023). _R: A Language and Environment for Statistical Computing_. R Foundation for Statistical Computing, Vienna, Austria. https://www.R-project.org/

Reddy, D. V. S. S., Kowshik, K., Kishor, M. J., Chittaranajan, M., and Sravani, E. 2018. Investigation of chitosan bio-polymer effect on the geotechnical properties of an expansive soil. In Proceedings of International Conference on Recent Trends in Engineering Materials, Management and Sciences, ICRTEMMS-2018 (pp. 25–27).

Ren, J., Tong, J., Li, P., Huang, X., Dong, P., and Ren, M. 2021. Chitosan is an effective inhibitor against potato dry rot caused by *Fusarium oxysporum*. Physiological and Molecular Plant Pathology, 113, 101601. 10.1016/j.pmpp.2021.101601

Rhoades, J. D. 1982. Cation exchange capacity. Methods of soil analysis: Part 2 chemical and microbiological properties, 9, 149–157. 10.2134/agronmonogr9.2.2ed.c8

Sawaguchi, A., Ono, S., Oomura, M., Inami, K., Kumeta, Y., Honda, K., Sameshima-Saito, R., Sakamoto, K., Ando, A., and Saito, A. 2015. Chitosan degradation and associated changes in bacterial community structures in two contrasting soils. Soil Science and Plant Nutrition, 61(3), 471–480. 10.1080/00380768.2014.1003965

Stöveken, J., Singh, R., Kolkenbrock, S., Zakrzewski, M., Wibberg, D., Eikmeyer, F. G., Pühler, A., Schlüter, A., and Moerschbacher, B. M. 2015. Successful heterologous expression of a novel chitinase identified by sequence analyses of the metagenome from a chitin-enriched soil sample. Journal of Biotechnology, 201, 60–68. 10.1016/j.jbiotec.2014.09.010

Suarez-Fernandez, M., Marhuenda-Egea, F. C., Lopez-Moya, F., Arnao, L. B., Cabrera-Escribano, F., Nueda, M. J., Gunsé, B and Lopez-Llorca, L. V. 2020. Chitosan Induces Plant Hormones and Defenses in Tomato Root Exudates. Front. Plant Sci. 11, 572087. 10.3389/fpls.2020.572087

Trolinger J.C., McGovern R.J., Elmer W.H., Rechcigl N.A. and Shoemaker C.M. 2017. Diseases of Chrysanthemum. In: R. McGovern, and W. Elmer (Eds.) Handbook of Florists’ Crops Diseases. Handbook of Plant Disease Management. Cham, Switzerland: Springer, pp. 439–491. 10.1002/ndr2.12013

Turan, V. 2019. Confident performance of chitosan and pistachio shell biochar on reducing Ni bioavailability in soil and plant plus improved the soil enzymatic activities, antioxidant defence system and nutritional quality of lettuce. Ecotoxicology and Environmental Safety, 183, 109594. 10.1016/j.ecoenv.2019.109594

Wagg, C., Schlaeppi, K., Banerjee, S., Kuramae, E. E., and van der Heijden, M. G. (2019). Fungal-bacterial diversity and microbiome complexity predict ecosystem functioning. Nature communications, 10(1), 4841. 10.1038/s41467-019-12798-y

Walkley, A., and Black, I. A. 1934. An examination of the Degtjareff method for determining soil organic matter, and a proposed modification of the chromic acid titration method. Soil science, 37(1), 29–38.

Wood DE., Lu J. and Langmead B. (2019). Improved metagenomic analysis with Kraken 2. Genome biology, 20(1), 257.

Wood DE. and Salzberg SL. 2014. Kraken: ultrafast metagenomic sequence classification using exact alignments. Genome Biology, 15(3), R46.

Zavala-González, E. A., Lopez-Moya, F., Aranda-Martinez, A., Cruz-Valerio, M., Lopez-Llorca, L. V., and Ramírez-Lepe, M. 2016. Tolerance to chitosan by *Trichoderma* species is associated with low membrane fluidity. Journal of Basic Microbiology, 56(7), 792–800. 10.1002/jobm.201500758

Zhan, J., Qin, Y., Gao, K., Fan, Z., Wang, L., Xing, R., Liu, S., and Li, P. 2021. Efficacy of a chitin-based water-soluble derivative in inducing *Purpureocillium lilacinum* against nematode disease (*Meloidogyne incognita*). International Journal of Molecular Sciences, 22(13), 6870. 10.3390/ijms22136870

