## Supplementary material for "Chitosan reduces naturally occurring plant pathogenic fungi and increases nematophagous fungus *Purpureocillium* under field soil conditions": Sup. Material

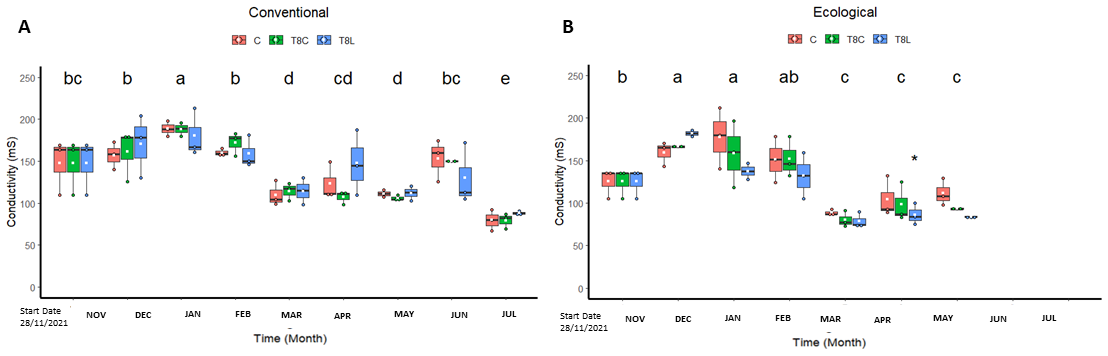

Figure S1: Effect of chitosan on field soil conductivity. Field soil was under conventional (A), or ecological (B) regimes. Treatments: Control (C, untreated), chitosan coacervate (T8C) and chitosan solution (T8L). Low-case letters show significant differences (p-value<0.05) with time.

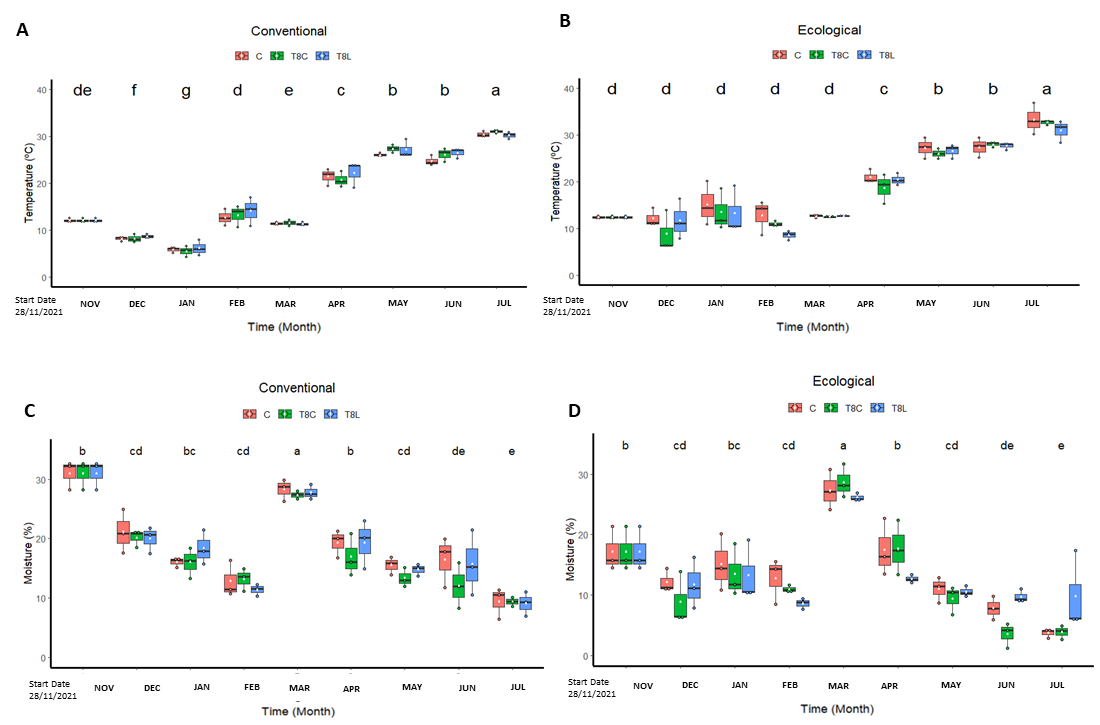

Figure S2: Effect of chitosan on field soil physical properties. Soil temperature A), B) and moisture C), D). Soil was field under conventional (A, C), or ecological (B) regimes. Treatments: Field (C, untreated), chitosan coacervates (T8C) and chitosan solution (T8L). Low-case letters show significant differences between the different times. Level of significant differences p-value<0.05.

Precipitation (mm)

Temperature (ºC)

| Time (month) | Precipitation |
| --- | --- |
| Noviembre | 9.69 |
| Diciembre | 0.852 |
| Enero | 2.068 |
| Febrero | 3.295 |
| Marzo | 85.9725 |
| Abril | 17.842 |
| Mayo | 21.5 |
| Junio | 0.4 |
| Julio | 1.52 |

|  | Temperature |
| --- | --- |
| Noviembre | 12.5003 |
| Diciembre | 12.1458 |
| Enero | 9.09452 |
| Febrero | 11.7589 |
| Marzo | 10.95 |
| Abril | 13.692 |
| Mayo | 19.3097 |
| Junio | 24.4937 |
| Julio | 26.4619 |

Figure S3: Monthly average precipitation and temperature in Pedralba (Valencian community, E, Spain).

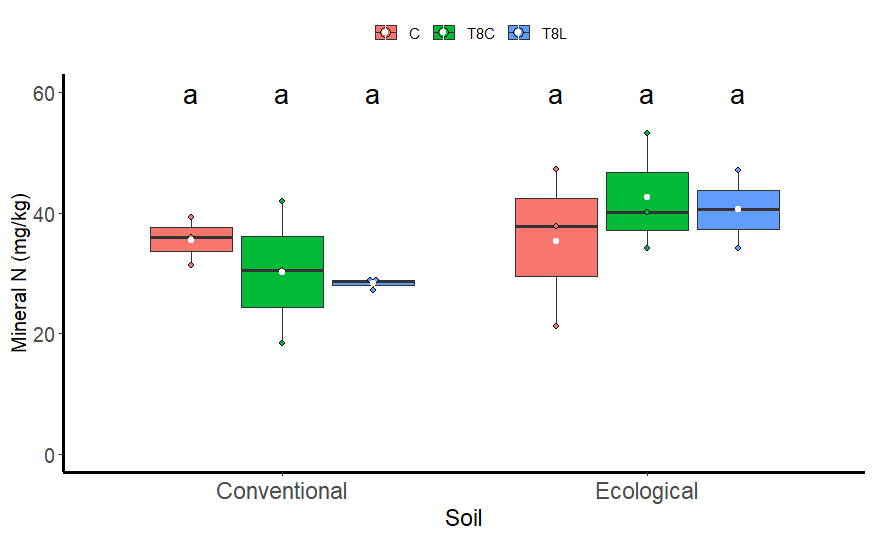

Figure S4: Effect of chitosan management on field soil mineral nitrogen content. Treatments: Control (C), chitosan coacervate (T8C) and chitosan solution (T8L).

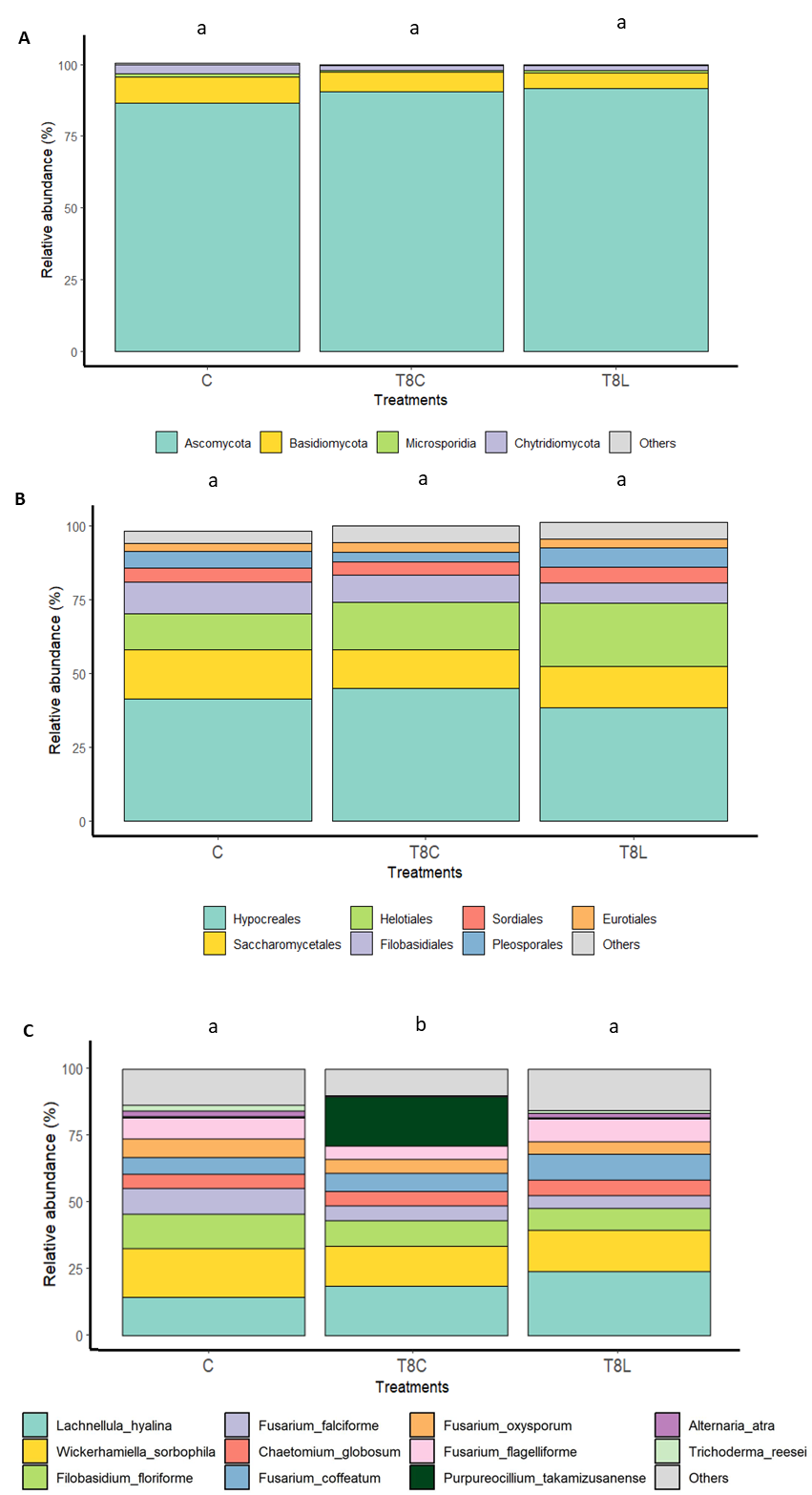

Figure S5: Effect of chitosan on field soil (Ecological Regime) Fungal microbiome (ITS primers). Phyla A), Orders B) and Species. Treatments: Field (C, untreated), chitosan coacervate (T8C) and chitosan solution (T8L). Different letters indicate significant differences (p-value < 0.05).

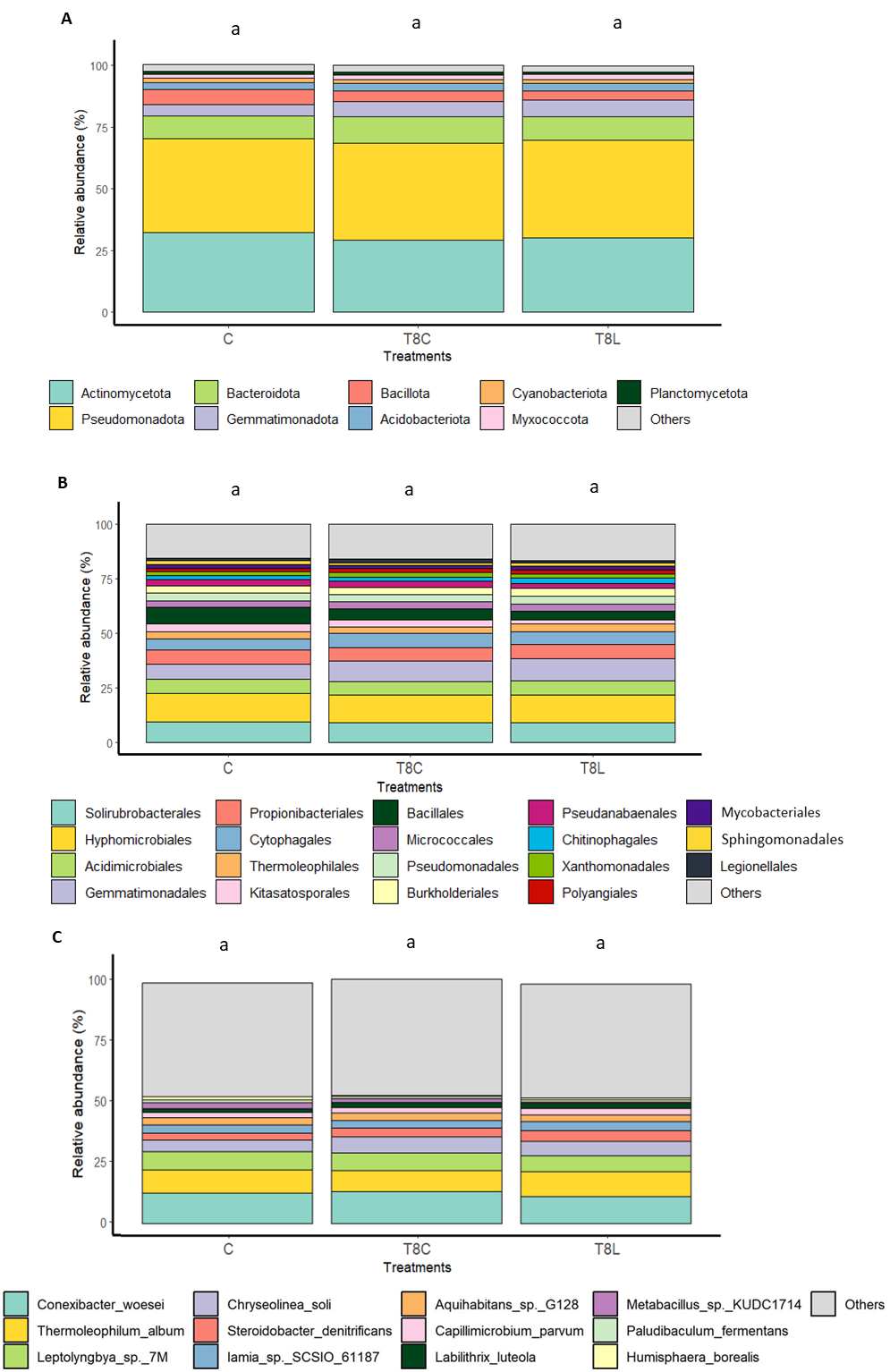

Figure S6: Effect of chitosan on field soil (ecological regime) Bacterial microbiome (V1-V2 primers). Phyla A), Orders B) and Species. Treatments: Field (C, untreated), chitosan coacervates (T8C) and chitosan solution (T8L). Different letters indicate significant differences (p-value < 0.05).

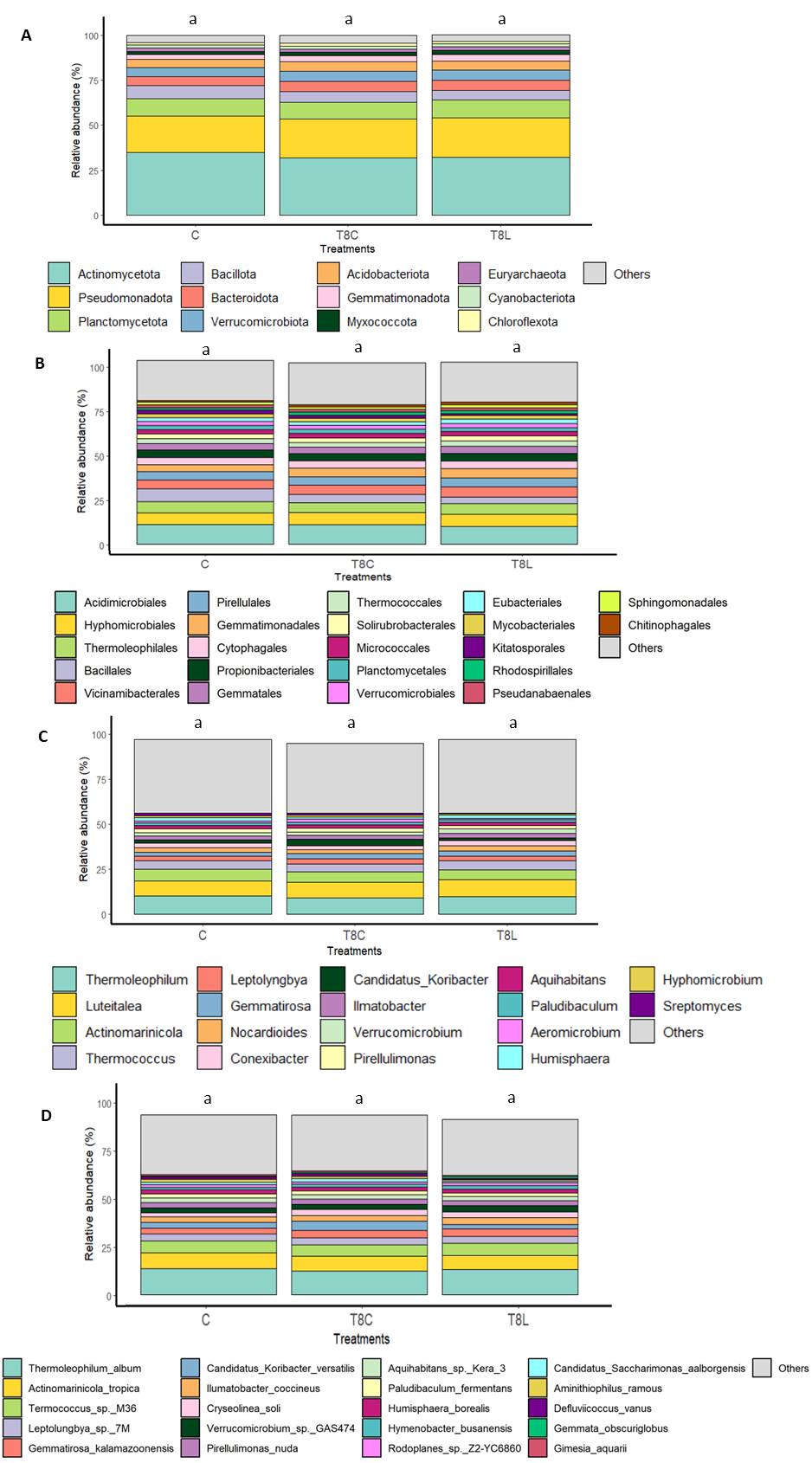

Figure S7: Effect of chitosan on soil (ecological regime) Bacterial microbiome. Phyla (A), Orders (B), Genera (C) and Species (D) (V3-V4 primers). Treatments: Control (C, untreated), chitosan coacervates (T8C) and chitosan solution (T8L). Letters show significant differences (p.value < 0.05).
